# Determination of the Two-Component Systems regulatory network reveals core and accessory regulations across *Pseudomonas aeruginosa* lineages

**DOI:** 10.1101/2021.07.23.453361

**Authors:** Julian Trouillon, Lionel Imbert, Anne-Marie Villard, Thierry Vernet, Ina Attrée, Sylvie Elsen

**Affiliations:** Université Grenoble Alpes, CNRS, CEA, IBS UMR 5075, Team Bacterial Pathogenesis and Cellular Responses, 38044, Grenoble, France; Université Grenoble Alpes, CNRS, CEA, IBS UMR 5075, 38044, Grenoble, France; Université Grenoble Alpes, CNRS, CEA, EMBL, ISBG UAR 3518, 38044, Grenoble, France

## Abstract

*Pseudomonas aeruginosa* possesses one of the most complex bacterial regulatory networks, which largely contributes to its success as a human opportunistic pathogen. However, most of its transcription factors (TFs) are still uncharacterized and the potential intra-species variability in regulatory networks has been mostly ignored so far. Here, to provide a first global view of the two-component systems (TCSs) regulatory network in *P. aeruginosa*, we produced and purified all DNA-binding TCS response regulators (RRs) and used DAP-seq to map the genome-wide binding sites of these 55 TFs across the three major *P. aeruginosa* lineages. The resulting networks encompass about 40% of all genes in each strain and contain numerous new important regulatory interactions across most major physiological processes, including virulence and antibiotic resistance. Strikingly, the comparison between the three representative strains shows that about half of the detected targets are specific to only one or two of the tested strains, revealing a previously unknown large functional diversity of TFs within a single species. Three main mechanisms were found to drive this diversity, including differences in accessory genome content, as exemplified by the strain-specific plasmid in the IHMA87 outlier strain which harbors numerous binding sites of chromosomally-encoded RRs. Additionally, most RRs display potential auto-regulation or RR-RR cross-regulation, bringing to light the vast complexity of this network. Overall, we provide the first complete delineation of the TCS regulatory network in *P. aeruginosa* that will represent an important resource for future studies on this pathogen.

## INTRODUCTION

Transcription factors (TFs) are major actors in the regulation of gene expression. From their action on DNA and their interaction with RNA polymerase, other regulatory proteins or signal molecules, results the activation or repression of gene expression, thereby dictating cellular physiology (Mejía-Almonte et al., 2020). Therefore, the characterization of TFs and of their target genes constitutes a major goal across most fields of biological research. Recently, a new *in vitro* cistromic approach called DAP-seq was developed for the *in vitro* large-scale analysis of TFs binding sites (TFBSs) and notably allowed the characterization of hundreds of TFs in the plant model organism *Arabidopsis thaliana* due to its high scalability when used in combination with cell-free protein expression (Bartlett et al., 2017; O’Malley et al., 2016). Since then, DAP-seq has been used by us and others to analyze the TFBS landscapes of different bacterial regulators and showed high sensitivity, allowing the delineation of key regulatory features (Garber et al., 2018; Trouillon et al., 2020, 2021; Zhang et al., 2020). Although it is believed that transcription regulatory network are highly versatile and adaptable (Lozada-Chavez, 2006; Perez and Groisman, 2009; Price et al., 2008), as shown by *in silico* analyses revealing potential inter- and intra-species TF functional variability (Galardini et al., 2015), the vast majority of TF studies still focus on a single strain to assess a particular TF function.

The major human opportunistic pathogen *Pseudomonas aeruginosa* exhibits a high intrinsic resistance to antibiotics, a large arsenal of virulence factors and a great capacity to adapt to changing environments (Moradali et al., 2017). This latter characteristic relies in particular on a considerable number of TFs which constitute nearly 9% of all proteins (∼500) encoded by its genome (Rodrigue et al., 2000; Stover et al., 2000; Winsor et al., 2016). Among the different families of TFs are the response regulators (RRs) which together with their cognate signal-sensing histidine kinases (HKs) form the two-component regulatory systems (TCSs) (Mitrophanov and Groisman, 2008; Stock et al., 2000). TCSs are numerous in *P. aeruginosa*, about twice as abundant as in the model organism *Escherichia coli*, and those that have been studied were found to be pivotal for orchestrating important cellular processes such as antibiotic resistance or the so-called acute-to-chronic lifestyle switch (Valentini and Filloux, 2016). Recently, several attempts at the global characterization of TCSs relied on phenotypic characterization of loss-of-function mutants (Badal et al.; Gellatly et al., 2018; Kollaran et al., 2019; Wang et al., 2021a). However, while bringing new information on selected phenotypes, this approach does not globally address direct RR-target regulatory interactions. Among all RRs encoded in the PAO1 genome, 51 are predicted TFs and the binding sites of only 8 of them (GacA, AlgR, PhoB, PhoP, CzcR, GltR, DsbR and BfmR) have been determined at a genome scale so far (Bielecki et al., 2015; Fan et al., 2021; Huang et al., 2019; Xu et al., 2021; Yang et al., 2021; Yu et al., 2021). Consequently, even though the TCSs regulatory network is thought to be a major driver of *P. aeruginosa* adaptability to different environments, including during infection (Francis et al., 2017), only a small proportion of the genes directly regulated by RRs is known to date, representing a major knowledge gap in the understanding of this bacterium. This is notably due to the fact that RRs are only active *in vivo* in response to specific signals that are unknown in most cases, making it challenging to assess their role, let alone determining their entire regulons (Rajeev et al., 2020). Additionally, some RRs seem to be involved in cross-or co-regulations (Bielecki et al., 2015; Fan et al., 2021; Huang et al., 2019), which represent pivotal features of the network topology and thus call for a large-scale investigation of this phenomenon.

*P. aeruginosa* is a fast-evolving bacterium with a large intraspecies genetic diversity, gifting this pathogen with numerous different ways of establishing infection and resisting treatments. Three major phylogenetic lineages have been identified among *P. aeruginosa* strains (Freschi et al., 2015), each exhibiting many phenotypical specificities. Notably, strains from each lineage possess different major virulence factors, including the recently-discovered ExlBA two-partner secretion system in the PA7 lineage and the Type III Secretion System (T3SS) and different secreted toxins in the two others (Elsen et al., 2014; Freschi et al., 2019; Hauser, 2009; Huber et al., 2016). While both *P. aeruginosa* large regulatory network and genetic diversity act as pivotal features for its success as a pathogen, most of *P. aeruginosa* TFs are still uncharacterized and the intraspecies variability in this regulatory network has been mostly ignored so far.

In this study, we leveraged the large scalability and *in vitro* aspect of DAP-seq to investigate the entire TCSs regulatory network across the three major *P. aeruginosa* lineages. To that aim, we report 342 DAP-seq experiments over 55 RRs and three strains, PAO1, PA14 and IHMA879472 (IHMA87), more than doubling the number of TFs with a genome-wide binding profile in *P. aeruginosa*.

## RESULTS

### DAP-seq allows the investigation of the TCSs regulatory network

*P. aeruginosa* possesses one of the highest numbers of TCSs in bacteria, with 72 predicted RRs in PAO1, including 51 with DNA-binding domains (Figure 1A), which span five different RR families with specific domain architectures (Figure 1B) (Francis et al., 2017). Across the PAO1, PA14 and IHMA87 strains, which each represents one of the three main *P. aeruginosa* phylogenetic lineages (Freschi et al., 2015), there is a total of 55 unique DNA-binding RRs, 48 of which are present in all three strains (Figure 1C; Table S1). To investigate the *P. aeruginosa* TCSs regulatory network, we expressed all 55 RRs fused to a N-terminal polyhistidine tag in *E. coli*-derived cell-free extracts and used them for DAP-seq with the fragmented genomes of PAO1, PA14 and IHMA87 in duplicates, resulting in 342 independent DAP-seq experiments, including controls. Since RRs usually need to be phosphorylated to be active, we used acetyl phosphate for RR activation as previously done (Garber et al., 2018; Zhang et al., 2020), which can allow auto-phosphorylation of the RR receiver domain *in vitro* and thus alleviates the need of knowing the activating signal to identify RRs binding sites. We overall identified binding sites in all three strains for the vast majority (49) of the 55 RRs (Table S1, Dataset S1). Six RRs did not yield reproducible binding sites on the three genomes (Table S2), including RRs from the two smaller subfamilies (Figure 1D), the LytTR- and ActR-like (each composed of only one RR: AlgR and RoxR, respectively) and thus did not seem to be active in our *in vitro* conditions. As expected for TFBSs, reproducible peaks were highly enriched in intergenic regions (Figure 1E). In order to ensure correct inference of gene targets from binding site locations and thus generate the TCSs regulatory network, transcriptional units (TUs) and promoter regions were defined based on TFBS recovery performance in comparison to experimentally obtained transcription start sites (TSSs) in PA14 (Wurtzel et al., 2012) and known TFBSs relative position from RegulonDB (Santos-Zavaleta et al., 2019) to optimize the sensitivity-specificity trade-off in our results (Figure S1, Methods). Peaks allowing target inference were mostly found in intergenic regions and, to a lesser extent, at the 3’-end of TUs (Figure 1F), matching the usual position of promoters (Santos-Zavaleta et al., 2019; Wurtzel et al., 2012). Furthermore, peaks found within putative promoter regions with a previously defined TSS in PA14 (Wurtzel et al., 2012) were centrally enriched at the core promoter (in the first 50-100bp upstream of TSS), as expected for TFBSs (Figure 1G).

**Figure 1.**
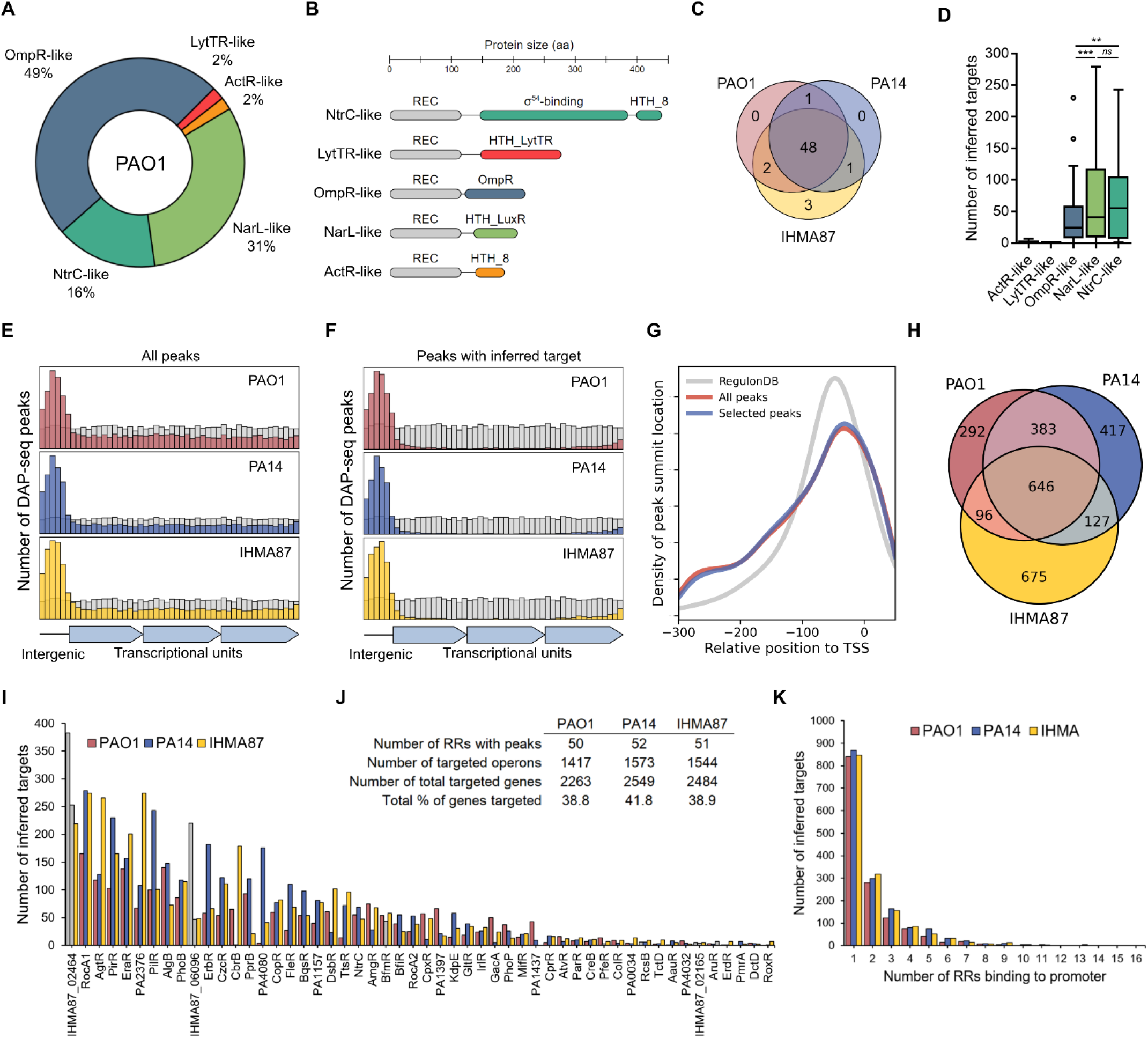
Overview of DAP-seq results. (A) Proportion of the five different RR subfamilies with a DNA-binding domain in *P. aeruginosa* PAO1. (B) Corresponding schematic representations of protein domain organizations. (C) Repartition of all DNA-binding RRs across the three strains PAO1, PA14 and IHMA87. (D) Number of inferred targets per RR families. Statistical significance was assessed using two-tailed t test (*p*-value < 0.01 [**] or 0.001 [***], *ns*: not significant). (E-F) Location of DAP-seq peak summits for all reproducible peaks (E) and all peaks that allowed target inference (F). Summits location in intergenic or inside of transcriptional units was assessed and rescaled to the genome’s intergenic/intragenic ratio (10.66-10.95% intergenic). Grey bins represent random distribution. (G) Density plot of peak summit location relative to known TSSs in PA14. The density of TFBSs location from RegulonDB is used as reference. (H) Repartition of RR inferred targets between strains. (I) Number of inferred targets for each RR with detected reproducible peaks. Grey bars represent non-physiological combinations (i.e. BfmR DAP-seq on the PA14 genome while PA14 does not possess *bfmR*). (J) Summary table of DAP-seq results. (K) Number of RR binding sites detected per target promoter.

Overall, this approach identified an average of 1,511 target TUs carrying a RR binding site in their promoter region per strain, encompassing 2,263-2,549 genes, or about 40% of all genes in each strain (Figure 1J, Dataset S2). To facilitate data exploration, we also provide a single table where all interactions are summarized and searchable by gene name (Table S2); further details such as fold enrichment and exact peak location can then be found in each RR DAP-seq results file (Dataset S1, Dataset S2). Interestingly, only 646 TUs, or 41-46% of all target TUs in each strain, exhibit a RR binding site in their promoter region in all three strains (Figure 1H), suggesting a high intra-species RR functional variability. As expected, PAO1 and PA14 show the highest overlap in identified target TUs. We also found large differences in number of targets between RRs, varying from seemingly very specialized RRs showing a very small number of targets (<10) to more global regulators with up to 285 inferred targets on average (Figure 1I). Additionally, 40-45% of targets were found to have more than one RR binding site on their promoter region (Figure 1K), suggesting a high proportion of co-regulation in the TCSs network. The analysis of the enriched DNA region allowed the determination of DNA binding motifs for most RRs (Figure 2A). The motifs often showed similarities between RRs with phylogenetically close DNA-binding domains, as seen for several groups of RRs (Figure 2B). Surprisingly, some RRs with seemingly different DNA-binding domains also showed very high similarity in DNA binding motifs, such as KdpE and CpxR (Figure 2C). Overall, DAP-seq allowed the near-complete determination of the TCSs regulatory network across the three tested strains and revealed several interesting key features of the network, as detailed below.

**Figure 2.**
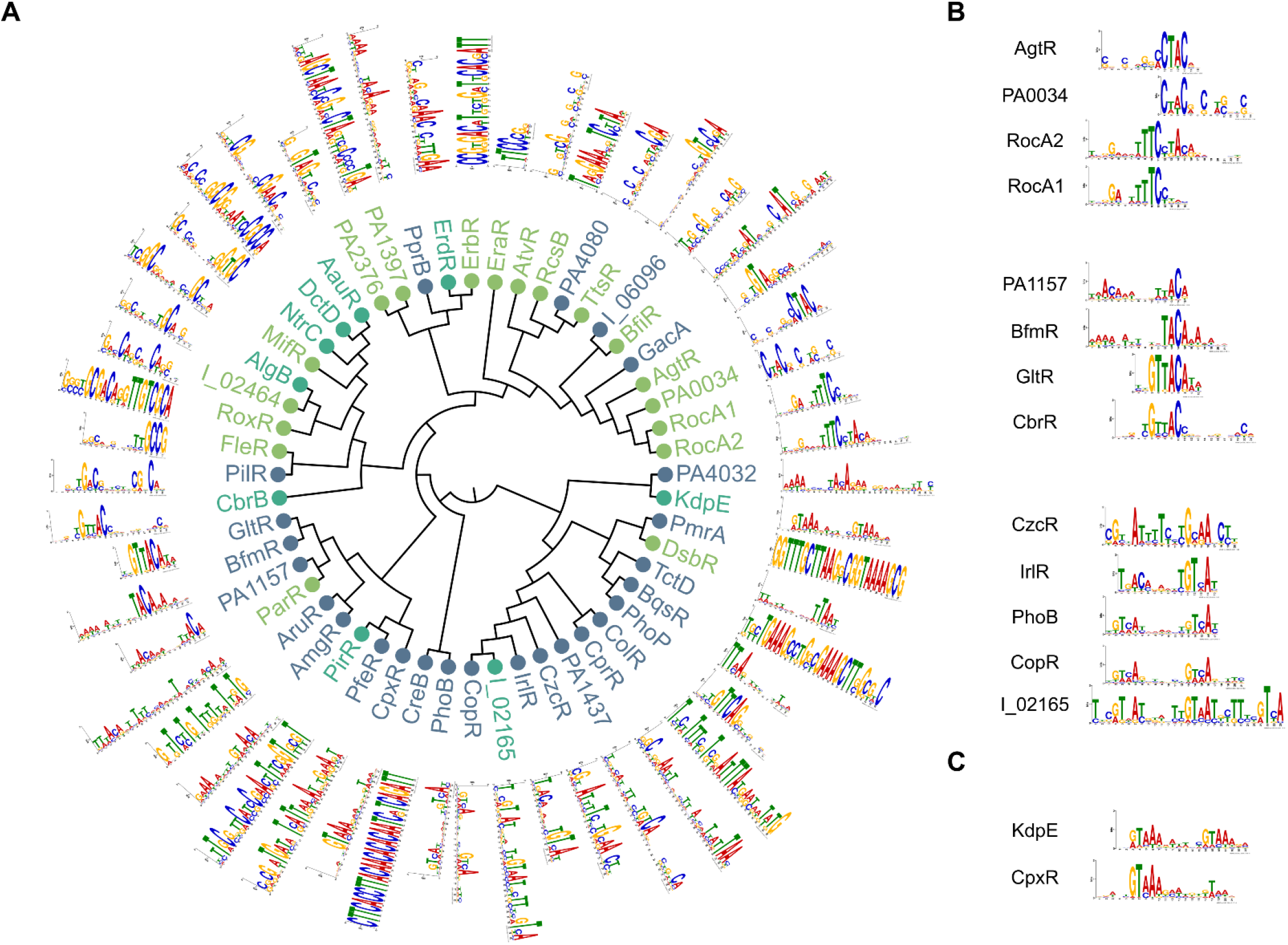
The global view of RRs DNA binding motifs. (A) Maximum-likelihood phylogenetic tree of 52 RR DNA-binding domains. The different OmpR, NarL and NtrC subfamilies are highlighted in blue, green and cyan, respectively. The DNA-binding motif found in peak regions using MEME-ChIP is shown for each RR. RR names with the “IHMA87_xx” format were shortened to “I_xx”. (B) Three groups of RRs that are neighbors in the tree and display similar DNA-binding motifs. (C) Two RRs with phylogenetically distant DNA-binding domains but sharing similar DNA-binding motifs.

To confront our datasets with the existing knowledge on *P. aeruginosa* RRs, we first compared it to known high-confidence targets, mostly from EMSA evidences. The comparison to this set extracted from 22 previous studies and composed of 59 known binding sites concerning 21 RRs showed that the vast majority were effectively retrieved, often resulting in very highly enriched peaks (Figure S2), further validating our approach. We then compared our results to the previously reported ChIP-seq studies on RRs for which we detected binding sites (PhoB, GacA, GltR, PhoP, CzcR, DsbR and BfmR) (Bielecki et al., 2015; Fan et al., 2021; Huang et al., 2019; Xu et al., 2021; Yang et al., 2021; Yu et al., 2021). Here again, there was a strong overlap in detected binding sites between the DAP-seq data and the six different ChIP-seq studies, as well as highly similar identified DNA motifs (Figure S3). The only exception was GacA; while our results revealed several new putative binding sites in addition to its two universally-recognized targets, *rsmY* and *rsmZ* (Brencic et al., 2009), only three targets overlapped with the previous ChIP-seq experiment, which however did not identify *rsmZ* as a GacA target (Figure S3C). Globally, our DAP-seq results confirmed known targets and expended the landscape of known RRs binding sites.

### The core TCSs regulatory network

The comparison of DAP-seq results between strains allowed the determination of the core TCSs regulatory network for the 48 conserved RRs, composed of 634 conserved RR-target interactions that were found in all three strains (Table S2). The clustering analysis of this core network resulted in the identification of 12 regulatory modules, each containing 1 to 7 RRs (Figure 3A). Notably, two larger modules including five (RocA1, RocA2, AgtR, TrsR and PA0034) and seven RRs (PirR, AmgR, KdpE, CpxR, PA1157 and GacA), showed high levels of gene target co-regulations across numerous biological processes. This analysis revealed both known (i.e. RocA1 and RocA2) and new (i.e. AmgR, PA1157, KdpE, CpxR and PirR) groups of RRs co-regulating common target genes. Some modules were composed of RRs with similar DNA motifs (Figure 2B–C), such as with RocA1, RocA2, AgtR and PA0034. These similarities in DNA motifs probably explain the larger proportion of shared targets between these TFs as they might share binding sites on the corresponding promoters. Numerous gene targets encoding key virulence factors were found to be part of the core network. For example, key operons encoding bacterial motility appendages such as the type IV pili and flagellum emerge in several clusters (Figure S4A-B). Moreover, several key regulatory genes were also targeted, including the two non-coding RNAs RsmY and RsmZ as well as their cognate RNA-binding protein RsmA (Figure 3A), all three being responsible for the regulatory switch between motile and sessile lifestyles (Moradali et al., 2017), and thus representing a pivotal node in *P. aeruginosa* regulatory network. Strikingly, six RRs were found to bind to at least one promoter of these three genes in all strains (Figure S4C), including the known regulator of RsmY and RsmZ, GacA.

**Figure 3.**
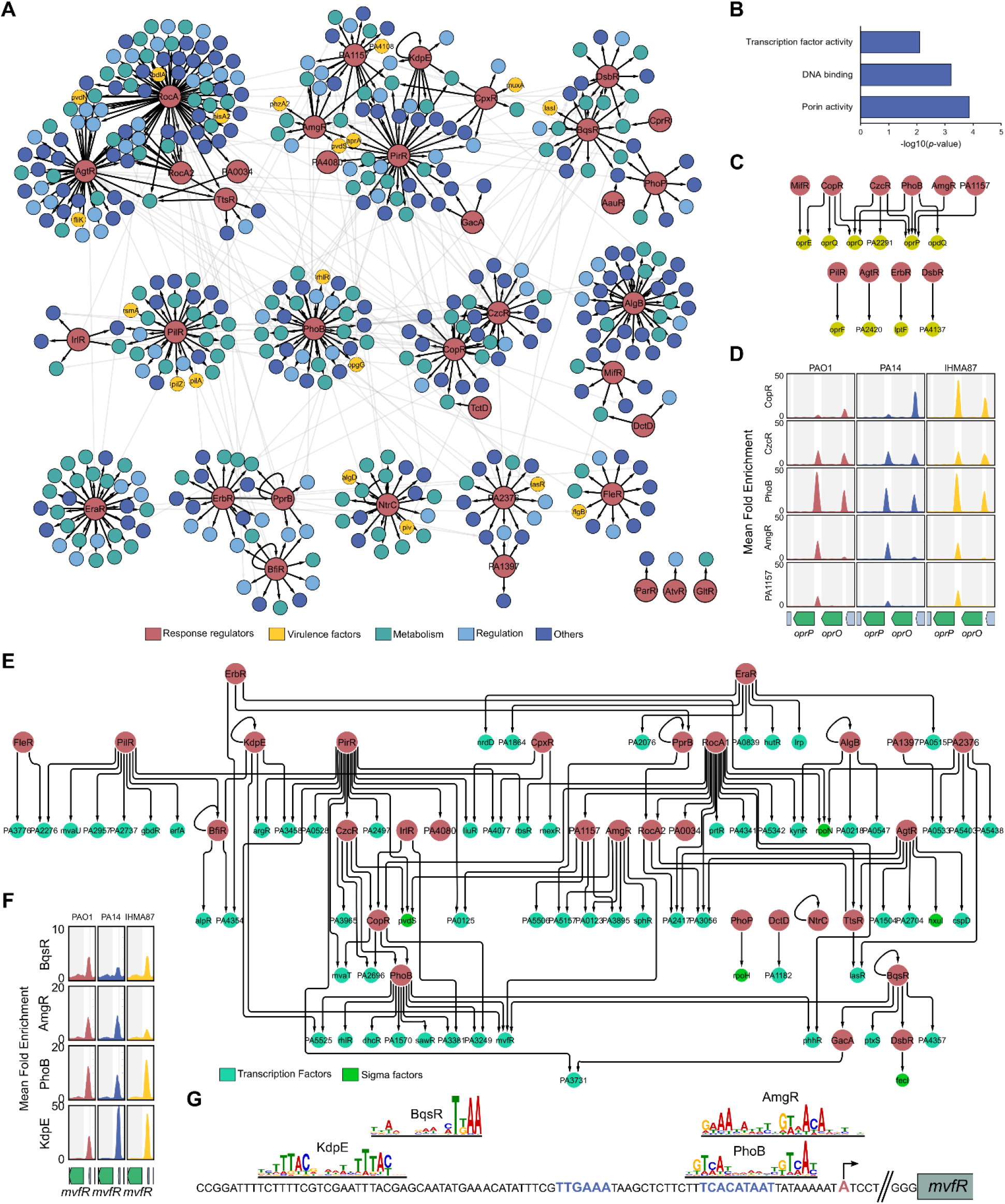
The core TCSs regulatory network. (A) Graph diagram of the core TCSs regulatory network. Cluster identification was performed using GLay (Su et al., 2010). Each cluster is shown as an independent module of the graph with black edges. Inter-cluster interactions are shown as light grey edges. RRs are colored in red and target TUs are colored depending on groups of COG predicted functions. The “Virulence factor” category comprises genes annotated as such in the Pseudomonas Database (Winsor et al., 2016). For operons, the name of the first gene is given. (B) Molecular Function GO Term enrichment analysis of the target genes of the core TCSs regulatory network. All GO terms with *p*-value < 0.01 are shown. (C) Graph diagram of all core interactions involving target genes annotated with the GO Term ‘Porin activity’. (D) Enrichment coverage tracks of DAP-seq against negative controls are shown for the five RRs with binding sites on the promoters of *oprP* and/or *oprO* in all three genomes. (E) Graph diagram of all core interactions involving target genes with a DNA-binding domain (Winsor et al., 2016). Each target node represents a TF-encoding gene found in target TUs from the core regulatory network. (F) Enrichment coverage tracks of DAP-seq in *mvfR* region. (G) Schematic view of *mvfR* promoter. RR motifs are shown at the location where they were identified by MEME. The TSS position experimentally determined in PA14 is shown in red and by a black arrow, predicted −10 and −35 boxes are in blue.

The functional enrichment analysis of the core network pointed out to two main overrepresented molecular functions among target genes: “porin activity” and “DNA-binding” (Figure 3B). Indeed, conserved RR binding sites were found in the promoter of 10 out of the 27 genes annotated with porin activity (Figure 3C). Two of these genes, encoding the OprO and OprP porins, were found with the highest numbers of RRs binding to their promoter regions in all three strains (3 and 5, respectively), suggesting the existence of an important regulatory node on these two neighboring genes (Figure 3D). While it was shown that PhoB regulates both genes (Bielecki et al., 2015), our results show that four additional RRs seem to be involved in the regulation of one or two of these porins. These include CzcR and CopR which share similar DNA-binding motifs with PhoB (Figure 2B), suggesting that they either compete for binding at these promoters or act under different conditions. Interestingly, OprO and OprP participate in phosphate uptake in *P. aeruginosa* (Chevalier et al., 2017) and while TCSs rely on phosphorylation for signal transduction and RRs activation, this result suggests the existence of several regulatory feedback loops from RRs onto phosphate uptake, which are probably important for correct TCS regulatory response (Klein et al., 2007).

The second and third most enriched molecular functions in target genes of the core TCSs regulatory network were DNA-binding and transcription factor activity, respectively (Figure 3B). Indeed, 82 TF-encoding genes, including 17 RRs and five sigma factors, are conserved RR targets (Figure 3E). Numerous major TFs were found targeted by several RRs, including the two quorum-sensing regulators, RhlR and LasR (Figure S4D-E), and the virulence regulator MvfR (PqsR), which was the TF with the highest number of conserved RR binding sites in its promoter (Figure 3F). The manual investigation of RR binding sites location on *mvfR* promoter suggested two activating (KdpE and BqsR) and two repressing (PhoB and AmgR) interactions (Figure 3G) and illustrates the ability of DAP-seq to precisely delineate protein-DNA interactions. Another example was the binding of PilR to the promoter of the *erfA* gene, encoding the repressor of the ExlBA two-partner secretion system in the IHMA87/PA7-like lineage (Figure S4F) (Trouillon et al., 2020). Additionally, the *exlBA* operon itself was found to be a target of PilR in IHMA87 (Dataset S2). Interestingly, PilR is the known activator of type IV pili genes (Ishimoto and Lory, 1992) that promote ExlBA-dependent cytotoxicity through host cell-bacterial contact (Basso et al., 2017). These two cooperating virulence factors might thus be co-regulated by PilR, revealing a new function for this RR in the *exlBA*^+^ IHMA87/PA7-like lineage.

Overall, these results demonstrate that the core TCSs regulatory network seems to be highly enriched in regulatory functions. With both feed forward loops for signal amplification and a high number of regulated TFs, it appears that the core regulatory topology of the network is conserved and place RRs relatively high in the hierarchy of *P. aeruginosa* regulatory network. Such a conserved and robust set of interactions between TFs co- and cross-regulating each other might allow for overall conservation of network topology and stable grafting of new strain- or lineage-specific functions in the accessory network, as explored below.

### Response regulators show functional variability between strains of *P. aeruginosa*

The pool of genes with a RR binding site on their promoter regions seems to vary between the three tested strains (Figure 1H). When looking at the 48 RRs present in all three strains, there is a difference between the global pool of genes targeted by at least one RR, even if not the same, in all three strains (Figure 4A) and the pool of conserved RR-target interactions (Figure 4B). This means that some target genes (*n*=114) harbor RR binding sites in their promoter in all three strains, but from different RRs. One such example is the *fleQ* gene, encoding for the major c-di-GMP-sensing biofilm regulator (Baraquet et al., 2012). PirR was found to bind to *fleQ* promoter in PAO1 and PA14 while RocA1 and RocA2 bound to this promoter in IHMA87 (Figure 4C). This difference could be explained by the absence or presence of the respective RR DNA-binding motifs in the *fleQ* promoter across the three strains (Figure 4D), hinting at a potential large diversity of regulation for genes with divergent promoters in our dataset. In the light of these observed differences between strains, we investigated the conservation of binding events and found various levels of conservation across all RRs and strains (Figure 4E). Interestingly, about 50 and 70 % of genes targeted by RRs in only 1 or 2 strains respectively, are conserved in more strains than that (Figure 4F), revealing that the absence/presence of genes explains only partially the regulatory differences between strains, while others are probably due to differences at the promoter level, as seen for *fleQ* (Figure 4C–D). This result highlights the necessity of conducting comparative approaches, even between closely related strains.

**Figure 4.**
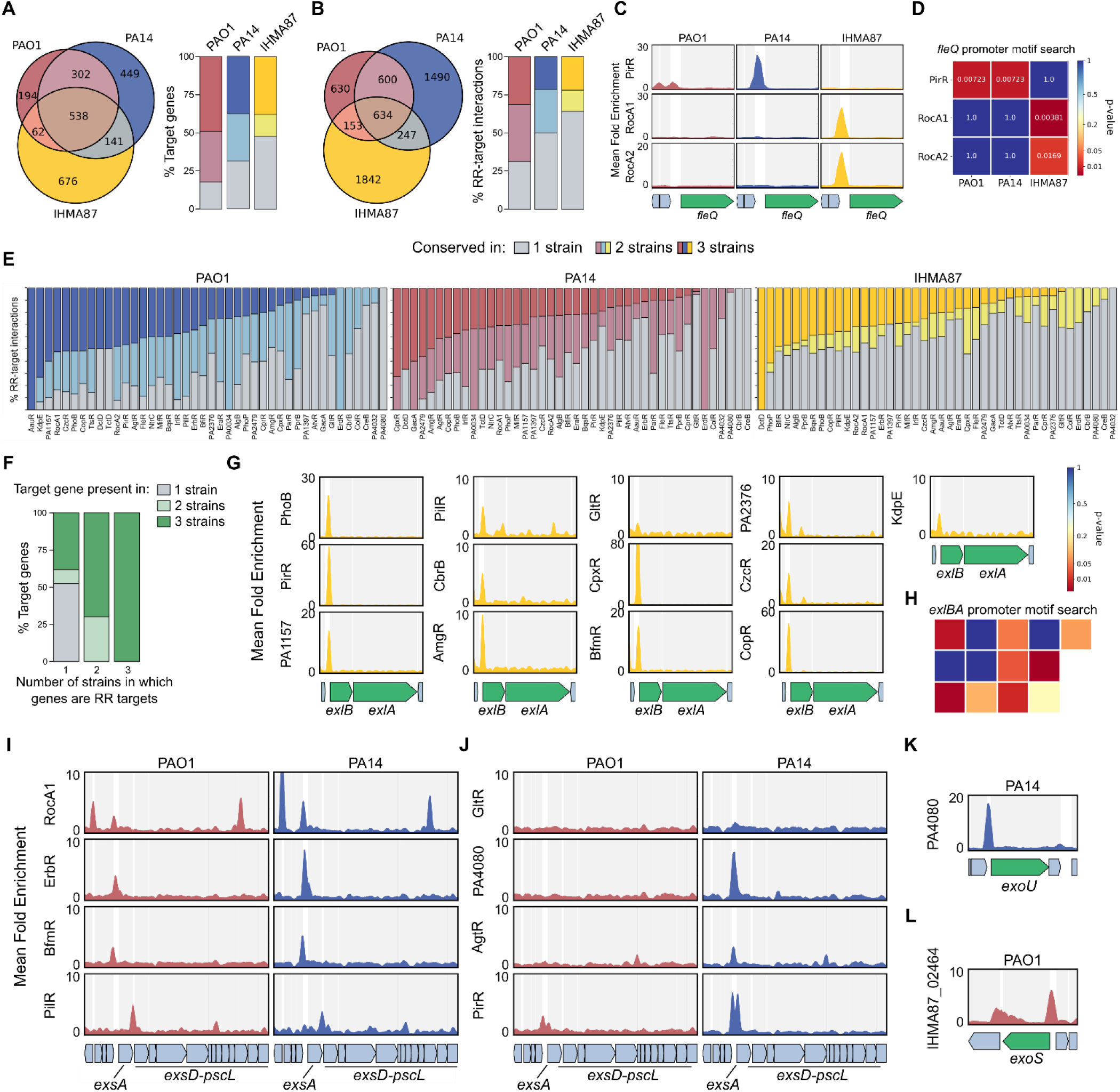
The accessory TCSs regulatory network. (A-B) Repartition of the total pool of gene targets (A) or RR-target interactions (B) between strains. (C) Enrichment coverage tracks of DAP-seq against negative controls for the three RRs with binding sites on the promoter of *fleQ* in all three genomes. (D) DNA-binding motif presence for the three RRs shown in (C) in *fleQ* promoter region in all three genomes, as shown as a heatmap of *p-value* obtained from motif searches using FIMO (Grant et al., 2011). (E) Proportion of RR-target interactions detected in one, two or three genomes for all genomes and conserved RRs with detected targets. RRs are sorted by number of targets found in three genomes. (F) Conservation of genes presence across the three strains for RR target genes, depending on the number of strains in which they are targeted. (G) Enrichment coverage tracks of DAP-seq against negative controls for the 13 RRs with binding sites on the promoter of *exlBA* in IHMA87. (H) DNA-binding motif presence for the 13 RRs shown in (G) in *exlBA* promoter region in IHMA87, as shown as a heatmap of *p-value* obtained from motif searches using FIMO (Grant et al., 2011). The RR order and repartition in the heatmap follow the organization of panel G. (I-L) Enrichment coverage tracks of DAP-seq against negative controls for the RRs with binding sites on T3SS-related genes in PAO1 and PA14, for conserved (I) or PA14-specific (J) T3SS-related RR-target interactions for shared genes, or for strain-specific genes in PAO1 (K) or PA14 (L).

Another source of regulatory differences between strains is the difference of RRs content itself. In the three strains tested here, seven RRs are present in only one or two strains (Figure 1C). As seen for PfeR, which is absent in PA14 but present in PAO1 and IHMA87, where it binds to the *pfeA* promoter (Dataset S2). Additionally, AlgB was found to bind to the *pfeA* promoter specifically in PA14, potentially representing a backup mechanism for the lost regulatory interaction (Dataset S2). Other such examples are the three IHMA87-specific RRs, especially IHMA87_02464 which harbors one of the highest number of inferred targets, with 219 in IHMA87 (Figure 1I). Among these, about 70% are genes also present in PAO1 and PA14, and IHMA87_02464 was found to be able to bind to the majority of them on the genomes of these two strains in our DAP-seq experiments (Dataset S2). Consequently, if this TF were to be acquired in one of these two strains, as common between bacteria through horizontal gene transfer (Price et al., 2008), it would already have numerous binding sites and regulatory targets.

A third mechanism of regulatory diversification stems from differences in target genes conservation. We found that 50% and 30% of genes targeted in one or two strains, respectively, were part of the accessory genome and absent from the genomes of the other strains (Figure 4F). Nearly half of these cases were found in IHMA87 (Figure 4A), reflecting its higher phylogenic distance compared to PAO1 and PA14. While *P. aeruginosa* is a highly diverse species with a core genome representing only 1.2% of all genes (Freschi et al., 2019), our results highlight how far we still are from apprehending TFs functions across bacterial species. One important difference often stressed between the three lineages is the mutual exclusion between the T3SS and ExlBA secretion systems, defining the pathogenic strategy of the given strain (Huber et al., 2016). Strikingly, we found 13 RRs binding to the *exlBA* promoter (Figure 4G), representing one of the most targeted promoters in this strain. While four of these binding peaks did not encompass the corresponding RR DNA motif (Figure 4H), nine of them did, revealing this promoter as a very highly connected regulatory node, especially since two additional TFs are already known to bind to it and to regulate *exlBA* (Berry et al., 2018; Trouillon et al., 2020). Furthermore, the genes encoding various T3SS components, notably the T3SS master regulator ExsA, were targeted by numerous RRs (Figure 4I–L). Four RRs were found to bind to the same locations in the major T3SS gene cluster in both PAO1 and PA14 (Figure 4I), with additional RRs binding there in PA14 (Figure 4J). Additionally, the two genes encoding the strain-specific toxins, which are found at different loci, *exoS* in PAO1 and *exoU* in PA14, also showed RR binding events on their promoters (Figure 4K–L). On its own, this T3SS-related example across these two strains shows all three types of regulatory differences, with differences in RR content (Figure 4L), target gene content (Figure 4K–L) and DNA-RR affinity (Figure 4J). Overall, in accordance with the recent predictions that the T3SS-related genes might be targeted by as many as 26 TFs in PAO1 (Wang et al., 2021b), we see that the regulation of these two major virulence factors seems highly connected, involving numerous TFs to allow for a precise and tightly controlled expression.

### The IHMA87 plasmid represents a hub of regulatory network integration events

One notable feature of the IHMA87 strain is that it carries a 185kb plasmid specific to that strain (Trouillon et al., 2020), named pIHMA87, which represents a great opportunity to explore regulatory plasticity and regulatory network grafting. Indeed, the acquisition of this plasmid by IHMA87, bringing 200 new genes, potentially required the wiring of some of these genes to the pre-existing IHMA87 regulatory network, as often predicted for horizontally acquired genes (Price et al., 2008). Strikingly, we found RR binding sites on the promoter regions of 73 of the pIHMA87 genes, involving 36 RRs (Figure 5A–B), revealing a high level of regulatory grafting to the regulatory network. Some RRs notably exhibited numerous binding sites encompassing their DNA motif on pIHMA87 (Figure 5C), showing the potential immediate addition of the corresponding genes to their regulons due to the presence of their binding sites on the acquired plasmid. While the vast majority of pIHMA87 genes are completely uncharacterized and often code for proteins with poor functional annotations, it is interesting to see that the target gene with the highest number (9) of RR binding sites in its promoter is *IHMA87_06166*, encodes a predicted type IV secretion system conjugation protein (Figure 5D). One of the main functions of this type IV secretion system is to mediate the transfer of plasmid DNA by conjugation (Cascales and Christie, 2003), suggesting that this key function in plasmid conjugation might have to be tightly regulated for the acquisition and conservation of the pIHMA87 plasmid.

**Figure 5.**
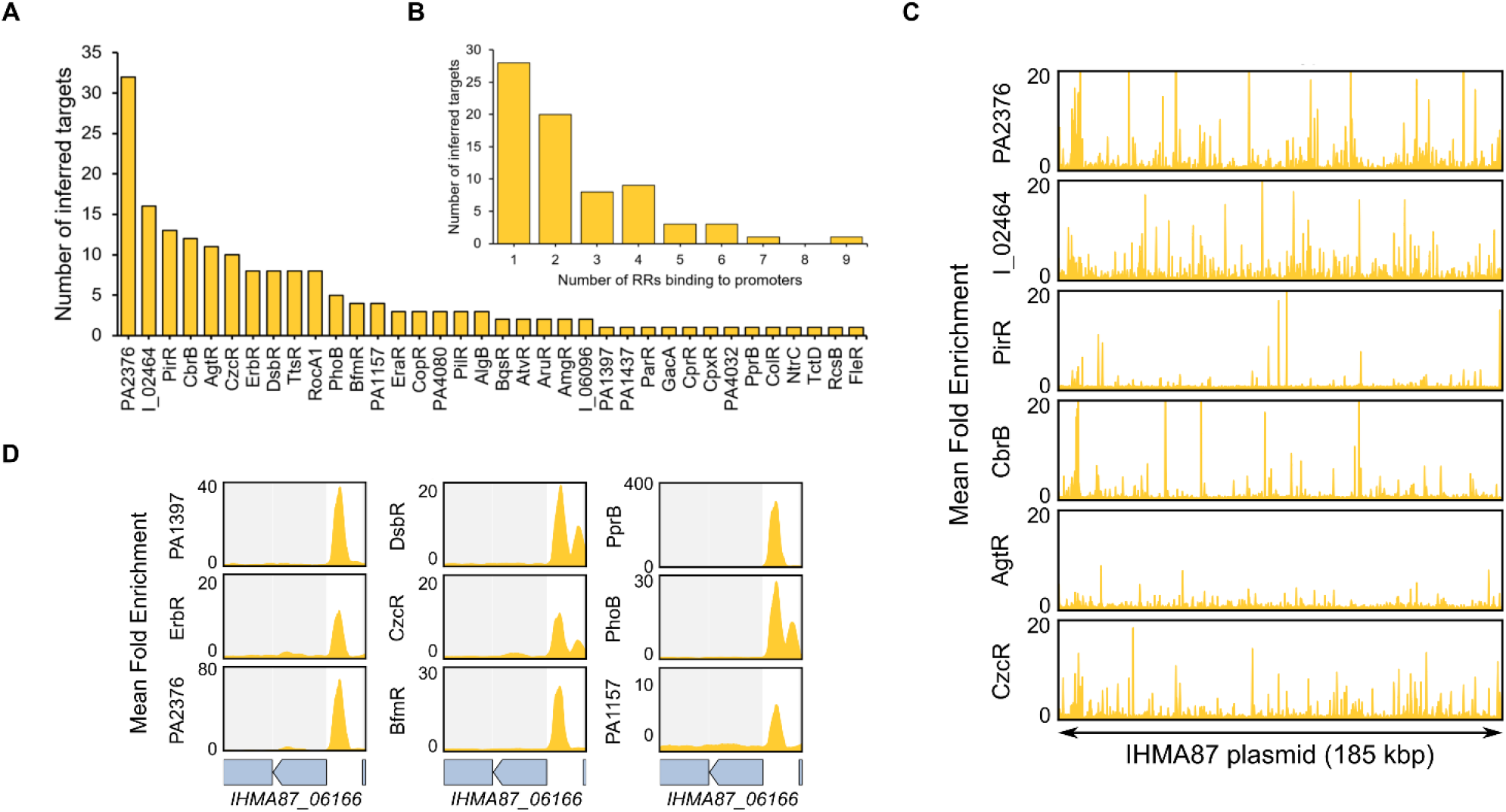
The integration of the pIHMA87 plasmid into the TCSs regulatory network. (A) Number of inferred targets on pIHMA87 per RR shown for all RRs with at least one plasmid binding site. RR names with the “IHMA87_xx” format were shortened to “I_xx”. (B) Number of RR binding sites detected per target promoter on pIHMA87. (C) Enrichment coverage tracks of DAP-seq against negative controls for the six RRs with >10 binding sites in promoter regions on pIHMA87. (D) Enrichment coverage tracks of DAP-seq against negative controls for the nine RRs with a binding site on the promoter of *IHMA87_06166* on pIHMA87.

### The response regulators network redundancy and complexity

The present analysis revealed two interesting features of the TCSs regulatory network: i) highly connected nodes representing target genes with numerous RR binding events on their promoter and ii) a high proportion of RR-RR cross-regulation. As exemplified for some genes above, we found that numerous target genes exhibited a high number of RR binding sites in their promoter, with 52 genes harboring >5 RR binding sites on average across the three strains (Table S2), going as high as 16 different RRs binding to one promoter. Among the most targeted genes are uncharacterized genes (*PA0123* and *PA3601*) but mostly well-characterized genes involved in bacterial survival and virulence (i.e. *flp*, encoding the type IVb pilin, the *nuoA-N* operon involved in the electron transport chain, *zipA*, encoding the ZipA cell division protein or the non-coding RNA RsmY) (Figure S5A). The gene encoding the alkaline protease AprA, a key secreted virulence factor important for immune evasion (Bardoel et al., 2012), stood out with the highest average number (13) of RRs binding sites (Figure S5B). The second and third most targeted genes code for the uncharacterized transcription factor PA0123 and the Xcp type II secretion system, involved in the secretion of several exoproteins including the ToxA and LasB toxins (Durand et al., 2003). All three corresponding promoters were found with >10 RR binding events on average, representing the most connected target nodes of the network. While PA0123 is considered uncharacterized, it was found to bind to the promoter of *lasR in vitro* and its overexpression induces strong cell growth inhibition (Longo et al., 2013), suggesting a potential important role for this transcription factor and calling for further characterization of its precise role. Overall, it appears that some genes requiring tight regulation exhibit complex promoter organization, allowing the potential coordination of an unusually high number of TFs acting on a single transcriptional unit.

One of the key observed features of the RR network was the high number of regulated TFs (Figure 3E), and particularly RRs. Indeed, numerous RRs were found to bind to the promoters of RR-encoding genes/operons, representing the RR cross-regulatory network (Table S2, Figure 6A). In total, we identified 149 such RR-RR regulatory interactions, including 26 that were found in all three strains (Figure 6B). Surprisingly, the proportion of RR-RR interactions conserved in all three strains was lower than the average of all inferred interactions (Figure 4), suggesting a more strain-specific need for these cross-regulation events. There was notably a high number of inferred auto-regulations, confirming few examples underlined in previous studies, such as for BqsR, PprB and PhoB (Bentzmann et al., 2012; Bielecki et al., 2015; Kreamer et al., 2015). Overall, some RRs seems to be more prone to either bind to many RR promoters (Figure 6C) or to have many RR binding sites on their promoter (Figure 6D). Among these, RocA1 targeted 12 RRs, including five in all three strains, placing this TF at the top of the TCS regulatory network hierarchy (Figure 6E). On the other hand, CzcR showed the highest number of RRs binding to its promoter region. Additionally, the contiguous *czcCBA* operon is in the top 10 most targeted transcriptional units globally (Table S2, Figure S5A), making the 512-bp *czcC*-*czcR* intergenic region the most targeted region, with a total of 22 RRs binding to it in at least one strain, including nine conserved in all three strains (Figure 6F). As for many such cases, while these regions are very similar between PAO1 and PA14, they share only ∼79% identity with the one in IHMA87, from which stems the vast majority of the non-conserved binding events. The *czcCBA* operon codes for an efflux pump responsible for the regulation of heavy metals concentration, which are the activating signal of the CzcRS TCS (Perron et al., 2004). Consequently, all RRs binding to this intergenic region could potentially regulate CzcR activity directly or indirectly, making CzcR a key downstream node of the RR network. Overall, these results highlight the high complexity of the TCS network, which contains numerous

**Figure 6.**
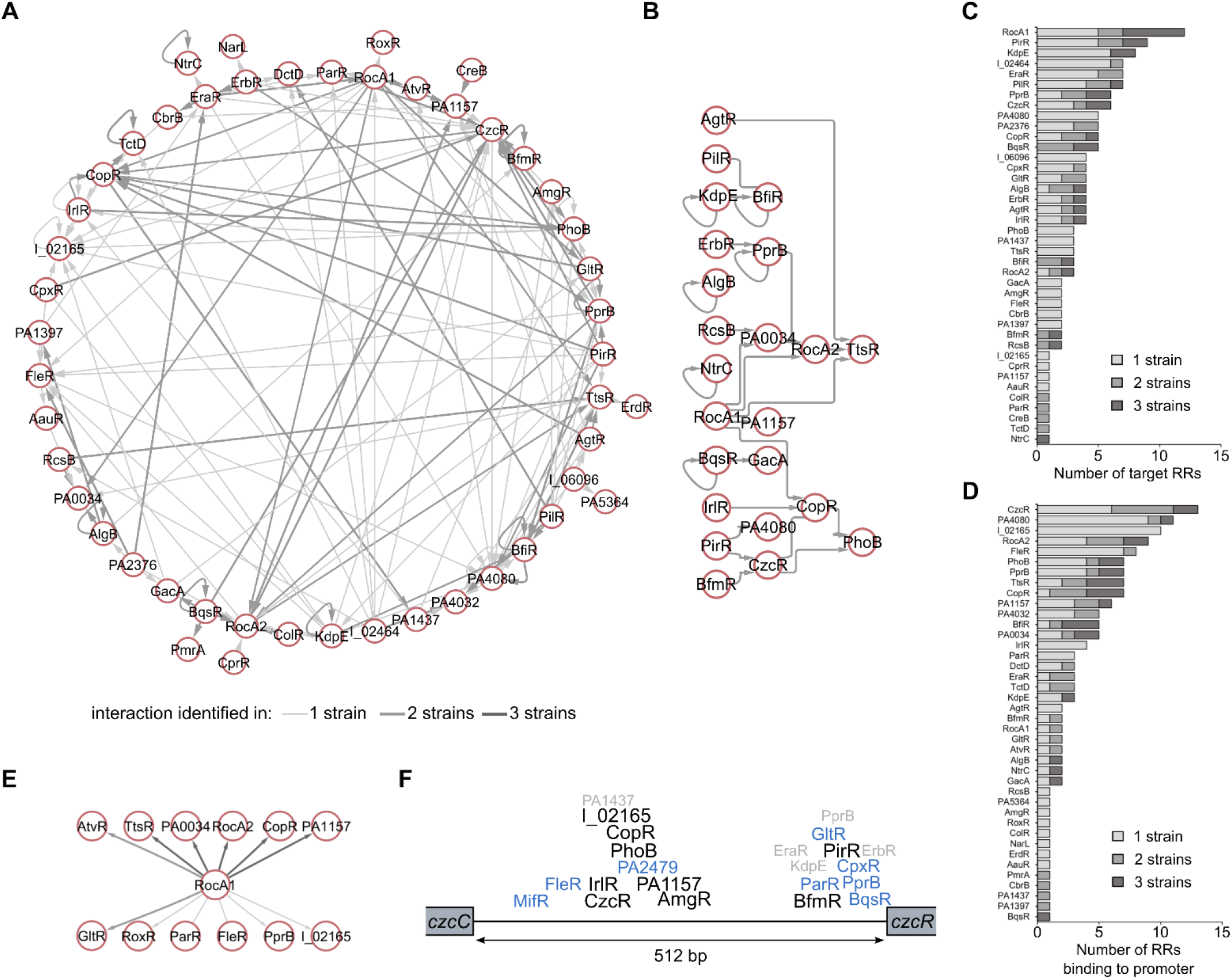
The RR cross-regulation network. (A) Graph diagram of all interactions involving RR target genes. Each node represents a DNA-binding RR found in target TUs from the TCS regulatory network. Edges thickness and darkness increase with the number of strains in which the interactions were identified. (B) Graph diagram of all RR-RR interactions found in all three strains. (C) Number of inferred RR targets for each RR. (C) Number of RRs binding to the promoters of each RR. (D) Graph diagram of all RR cross-regulation interactions involving RocA1 as source node. (E) Schematic representation of the binding events at the *czcC*-*czcR* intergenic region. The names of RRs that were found to bind to this region in one (grey), two (blue) or three strains (black) are indicated, centered at the average relative DAP-seq peak summit between strains. RR names with the “IHMA87_xx” format were shortened to “I_xx”.

RR cross-regulation allowing for potentiation or repression of signal transduction and thus fine-tuning of gene expression.

## DISCUSSION

Global health issues caused by bacterial pathogens are consistently increasing in scale and are now reaching alarming levels, mainly due to the development of antibiotic resistance (Blair et al., 2015) and host adaptation (Sheppard et al., 2018). Both mechanisms rely on acquisition or adaptation of regulatory functions, which mostly involve transcription factors. However, knowledge on bacterial TFs is still sparse; most of them are still uncharacterized even in the most studied organisms, and intra-species functional variability is hardly ever addressed, as the vast majority of studies are performed on a single well-established laboratory strain supposed to be representing its entire species. As a starting point to address the importance of this issue and to make our contribution to the needed characterization of *P. aeruginosa* TFs, we used DAP-seq to identify the binding sites of all RRs in three strains representing the three major *P. aeruginosa* lineages.

The main limitation when studying TCSs *in vivo* is the need of knowing the activating signal. As most signals are still unknown, this has considerably limited the ability of previous studies to characterize RRs globally (Rajeev et al., 2020). In this context, the *in vitro* aspect of DAP-seq and use of phosphate donor allowed us to circumvent this limitation and made the family-wide RR binding site investigation possible. While DAP-seq represents a powerful tool for the study of TFs, notably due to its higher scalability than ChIP-seq, it also comes with limitations. Indeed, as an *in vitro* assay working with individually purified proteins, DAP-seq might not fully picture the effect of TFs interacting with other molecules in the cell, such as TF-TF interactions, as previously seen for PhoB and TctD (Bielecki et al., 2015). More targeted work will be required to evaluate such interactions. A recent adaptation of the DAP-seq protocol was developed for the study of TF-TF interactions and their impact on DNA binding (Lai et al., 2020) and could notably allow such investigation. Additionally, in a similar way to what is observed for ChIP-seq using overexpressed TFs, some detected binding events might not completely reflect physiological interactions. For instance, while being present in all three strains, TtsR is an orphan RR in PAO1 and PA14, as its cognate HK, TtsS, is only present in the PA7/IHMA87 lineage (Cadoret et al., 2014). Consequently, similarly to the few non-physiological combinations performed here (i.e. BfmR DAP-seq on the PA14 genome while PA14 does not possess *bfmR*), the detected binding events in PAO1 and PA14 might not happen *in vivo* since TtsR might not be activated in these strains. On the other hand, DAP-seq identifies the full catalogue of binding abilities of a TF since, unlike ChIP-seq, it is not limited to one specific growth condition, and thus can potentially identify binding events that might occur in conditions for which the use of ChIP-seq is challenging, such as during infection.

Overall, our work represents the first inter-strain comparison of TF function of this scale and reveals numerous differences in TF binding landscapes between three strains of the same species. The observed differences arose from at least three main mechanisms: i) difference in TF contents, ii) difference in target gene content and iii) difference in target promoter sequences. Across the three strains tested here, the IHMA87 strain displayed the highest number of strain-specific interactions. This was expected as this strain is phylogenetically more distant from the other two and thus harbors many gene and promoter sequence differences. In addition, one major difference is the presence of the strain-specific plasmid in the IHMA87 strain, which contains 200 genes absent in the other strains. We found that 73 of these genes harbor a RR binding site on their promoter. This result exemplifies the striking capacity of acquired genes to graft to the existing regulatory network in the recipient bacterium. Indeed, previous *in silico* analyses showed that horizontally acquired genes are more prone to be regulated by multiple TFs and to be added to existing regulons, either through their concomitantly-acquired promoters that already carry the corresponding TF binding sites or through evolution of the binding sites after acquisition (Price et al., 2008). Although we cannot know which of the two happened for genes on IHMA87 plasmid, this mechanism plays a key role for phenotypic diversification and thus needs to be further investigated globally. In light of the huge genetic diversity across the wide *P. aeruginosa* species, and in fact across most bacterial species, it is obvious, from our results on only three strains, that TF functional diversity is omnipresent in bacteria and should be explored in much more details than what is currently done.

The functional characterization of TFs is a major goal that serves most biological fields of research. In *P. aeruginosa*, few major contributions towards this goal were recently published, mostly focused on virulence regulation (Huang et al., 2019; Wang et al., 2021b). Similarly, the present work significantly contributes to the characterization of *P. aeruginosa* TFs and constitutes an important resource for the community. While most RRs had been studied with targeted approaches, often identifying only one or few regulatory targets linked to a single studied phenotype, the complete direct regulons were still missing for ∼85% of the RRs. In most of these cases, our results confirm existing knowledge and complete it. Additionally, we provide binding profiles for 10 so far completely uncharacterized predicted RRs, such as PA1157 and PA4080. Overall, we report new RR binding sites in 1,511 promoters on average between the three strains. We believe that this dataset will serve the community and guide future studies for the delineation of regulatory and virulence key features of bacterial pathogens.

## METHODS

### Homolog determination, response regulators identification and phylogenetic analyses

All sequences and annotations were retrieved from the Pseudomonas Genome database v18.1 (Winsor et al., 2016). For inter-strains comparisons, homolog determination was performed by Reciprocal Best Blast Hit (RBBH) analysis (Cock et al., 2015) on the European Galaxy server (Jalili et al., 2020) using all protein sequences from *P. aeruginosa* PAO1, PA14 and IHMA87 strains with minimum percentage alignment coverages of 90 and sequence identities of 50. Functional classification of predicted RRs with DNA-binding domains were obtained from PseudoCAP (Brinkman et al., 2000), the P2TF database (Ortet et al., 2012) and the Pseudomonas Genome database (Winsor et al., 2016), and manually pooled and curated. All candidate RRs were then verified by protein classification using Interpro 73.0 (Blum et al., 2021) and candidates with confirmed identification of both a RR receiver domain (REC) and a DNA-binding domain were selected, resulting in the final list of 55 RRs across the three strains (Table S1). Pfam annotations from the Interpro search were used for DNA-binding domain sequence determination for all RRs, which were then used to generate a multiple alignment using MUSCLE (Edgar, 2004). The resulting alignment was used to build a maximum-likelihood phylogenetic tree using MEGA X with 100 bootstraps which was visualized and annotated using iTOL v5 (Kumar et al., 2018; Letunic and Bork, 2019).

### Plasmids and genetic manipulations

Primers are listed in Table S3. For production of recombinant proteins, the 55 gene sequences were amplified by PCR using genomic DNA as matrix and appropriate primer pairs and then integrated by Sequence- and Ligation-Independent Cloning (SLIC) (Li and Elledge, 2007) in pIVEX2.4d (Martin et al., 2001) cut with *Not*I-*Sal*I. PAO1 genomic DNA was used as template for the 51 RRs found in that strain, IHMA87 (3) and PA14 (1) DNA were used for the 4 remaining ones. All plasmids were transformed into competent TOP10 *E. coli* cells (ThermoFisher Scientific) and then checked by sequencing (Eurofins).

### Cell-free protein expression

Cell-free expression was performed for all constructs after Magnesium concentration optimization. The 55 RRs were expressed as previously described (Imbert et al., 2021) in a final volume of 100 μL for each RR during 2 h at 23°C under gentle agitation, with a batch mode configuration. A total of 16 μg.ml^−1^ of RR template DNA in pIVEX2.4d were added to a reaction mixture containing 1 mM of each of the 20 amino acids, 0.8 mM rNTPs (guanosine-, uridine-, and cytidine-5′-triphosphate ribonucleotides), 1.2 mM adenosine-5′-triphosphate, 55 mM HEPES, pH 7.5, 68 μM folinic acid, 0.64 mM cyclic adenosine monophosphate, 3.4 mM dithiothreitol (DTT), 27.5 mM ammonium acetate, 2 mM spermidine, 80 mM creatine phosphate, 208 mM potassium glutamate, 14 mM magnesium acetate, 250 μg.ml^−1^ creatine kinase, 27 μg.ml^−1^ T7 RNA polymerase, 0.175 μg.ml^−1^ tRNA and 40% S30 *E. coli* bacterial extract. Protein extracts were then clarified by centrifugation at 14,000 g for 20 min at 10°C and the resulting supernatants were used for DAP-seq.

### DAP-seq experimental procedure

Fragmented genomic DNA libraries were prepared exactly as previously described (Trouillon et al., 2020) using the purified genomic DNA of *P. aeruginosa* PAO1, PA14 or IHMA87 strains. DAP-seq was performed as previously described (Trouillon et al., 2020) with some modifications. Briefly, 100 μl of cell-free soluble protein extracts were diluted in 100 μl of Binding Buffer (sterile PBS supplemented with 10mM MgCl2, 0.01% Tween 20 and 50μM acetyl phosphate) containing 10 μl of pre-washed (3X) magnetic cobalt beads (Dynabeads His-Tag Isolation and Pulldown - Invitrogen) for a final volume of 200 μl in 96-well plates and incubated for 40 min at room temperature on a rotating wheel. The bead–protein complexes were then washed six times in 200 μl of Binding Buffer, including a transfer to a new 96-well plate before the last wash, and then resuspended in 80 μl of Binding Buffer containing 50 ng of adaptor-ligated gDNA libraries and further incubated on a rotating wheel at room temperature for 1 h. The bead–protein–DNA complexes were then washed six times in 200 μl of Binding Buffer, including a transfer to a new 96-well plate before the last wash. Beads were then resuspended in 30 μl of sterile 10 mM Tris-HCl pH 8.5, and incubated for 10 min at 98°C for elution. After incubation, samples were placed on ice for 5 min, beads were then magnetically removed and the released DNA was used for PCR amplification as previously described (Bartlett et al., 2017) using a different indexed pair of primers for each sample. PCR products were pooled for sequencing using 5 μl of each sample in pools of up to 104 samples. Library pools were then purified using SPRIselect beads at a 1:1 ratio. The quality of each library pool was assessed using High Sensitivity DNA chips on an Agilent Bioanalyzer and additional bead purifications were performed in case of excess amounts of primer dimers. Negative control experiments were done using a pIVEX2.4d vector expressing an untagged GFP protein for each DAP-seq 96-well plate, genome and sequencing pool in duplicates. All DAP-seq experiments were performed in duplicates.

### Sequencing & primary data analysis

Sequencing was performed at the high-throughput sequencing core facility of I2BC (Centre de Recherche de Gif – http://www.i2bc.paris-saclay.fr) using an Illumina NextSeq500 instrument for a total of 4 High Output runs. An average of 2.6 million single-end 75-bp reads per sample were generated with >95% of reads uniquely aligning to the corresponding genome using Bowtie2 (Langmead and Salzberg, 2012). Peak calling was done using MACS2 (Zhang et al., 2008) on all uniquely aligned reads with a *p*-value threshold of 0.0001 for each duplicate against a pool of the two corresponding negative control samples. Only peaks found in both replicates were then selected using the Intersect tool from BEDTools (Quinlan and Hall, 2010) with a minimum overlap of 50% on each compared peaks. Peaks that passed the Intersect selection were then filtered by Irreproducible Discovery Rate (IDR) with a IDR threshold of 0.005 between replicates using IDR Galaxy version 2.0.3 on the European Galaxy server (Jalili et al., 2020; Li et al., 2011), yielding the final list of reproducible peaks.

For genomic coverage visualization, coverage bedgraph files obtained from MACS2 callpeak used with the --SPMR option were used to generate enrichment tracks using MACS2 bdgcmp in linear scale FE mode for each replicate against the corresponding control samples. Enrichment tracks were averaged between duplicates and used for data visualization.

For DNA motif discovery, the 100-bp central region of reproducible peaks from all three genomes were used for each RR using MEME-ChIP (Machanick and Bailey, 2011) with default settings. For DNA motif scanning, MEME-ChIP output files were used to find motifs in selected sequences using FIMO (Grant et al., 2011).

### Transcription regulatory network inference

For inference of gene targets from DAP-seq peak locations, since transcriptional units (TUs) and transcriptional start sites (TSSs) experimental annotations are only available for less than a third of all genes in PAO1 and PA14 (Gill et al., 2018; Wurtzel et al., 2012) and not at all in IHMA87, we optimized a common *in silico* approach to perform genome-wide definition of promoter regions in all three strains through i) TUs annotation and ii) promoter region sizing. The two corresponding parameters - the operon prediction score threshold and the size of upstream promoter regions - were modified over a range of 18 different combinations used for complete target inference over the full dataset and for assessment of TFBS recovery against the entire RegulonDB TFBS database (Figure S1) (Santos-Zavaleta et al., 2019). The best-performing parameter pair was then chosen as allowing the highest recovery of known TFBS (>95%) while keeping the number of predicted TUs close to PA14 experimental number (0.994:1 ratio). Consequently, TUs were defined by genome-wide operon prediction for the three PAO1, PA14 and IHMA87 strains using Operon-mapper (Taboada et al., 2018) with a minimum score threshold of 0.9. Promoter regions were then defined as the −400 bp to +20 bp region based on the translational start of the first gene of each transcription unit. Peaks with summit position in promoter regions were then identified using the Intersect tool from BEDTools (Quinlan and Hall, 2010) and assigned to the corresponding gene or operon for generation of the network and inference of regulatory interactions. In case of peaks found in overlapping promoter regions, the peak was assigned to the closest gene start, but additional overlapping regions are still reported in the final peak output files as additional columns on the peak row (Dataset S2), to allow for manual investigation.

### ChIP-seq data analysis

Peak lists from ChIP-seq experiments with PhoB, GacA, CzcR, GltR, PhoP, DsbR and BfmR were used to retrieve all peak summit locations (Bielecki et al., 2015; Fan et al., 2021; Huang et al., 2019; Xu et al., 2021; Yang et al., 2021; Yu et al., 2021). Peak summit positions were averaged between replicates and used to infer gene targets using the genome-wide promoter regions defined above with the Intersect tool from BEDTools (Quinlan and Hall, 2010) on the corresponding genomes: PAO1 (Fan et al., 2021; Huang et al., 2019; Xu et al., 2021; Yu et al., 2021) or PA14 (Bielecki et al., 2015; Yang et al., 2021). For DNA motif determination, peak regions were extended by 50bp on each side of the summits and retrieved using the GetFastaBed tool from BEDTools (Quinlan and Hall, 2010). The obtained sequences were then used for motif analysis using MEME-ChIP (Machanick and Bailey, 2011) with default settings, with the exception of the GltR, DsbR and PA14 PhoB ChIP-seq datasets for which minimum fold-change thresholds of 10, 5 and 5, respectively, had to be applied to peaks to obtain significantly enriched motifs, as also described in the original articles.

### Network and functional enrichment analysis

Network analyses were performed on Cytoscape 3.8 (Shannon et al., 2003), using either GLay clustering (Su et al., 2010) or yFiles Hierarchical clustering (Wiese et al., 2004). Functional annotations were retrieved from the Pseudomonas database (Winsor et al., 2016) and GO functional enrichment analyses were performed using DAVID v6.8 (Huang et al., 2007).

## Materials Availability

Plasmids generated in this study are available from the Lead Contact.

## Data availability

DAP-seq data files have been deposited on NCBI Gene Expression Omnibus (GEO) and can be accessed through GEO Series accession number GSE179001.

## Author contributions

Conceptualization, J.T. and S.E.; Methodology, J.T.; Investigation, J.T., L.I., A.M.V., T.V. and S.E.; Formal analysis and Visualization, J.T.; Writing – Original Draft, J.T. and S.E.; Writing – Review & Editing, J.T., L.I., A.M.V., T.V., I.A. and S.E.; Funding Acquisition, S.E. and I.A..

## Acknowledgments

We acknowledge the RoBioMol (hosted by the Pneumococcus Group-IBS) and the Cell-free platforms of the Grenoble Instruct-ERIC center (ISBG; UAR 3518 CNRS-CEA-UGA-EMBL) within the Grenoble Partnership for Structural Biology (PSB), supported by FRISBI (ANR-10-INBS-0005-02) and GRAL, financed within the University Grenoble Alpes graduate school (Ecoles Universitaires de Recherche) CBH-EUR-GS (ANR-17-EURE-0003). We thank Dr. Jérôme Boisbouvier and Dr. François Parcy for initial discussions on cell-free protein expressions and DAP-seq, respectively. We acknowledge the High-throughput sequencing facility of I2BC for its sequencing and bioinformatics expertise. This work was supported by the French National Research Agency in the framework of the “Investissements d’avenir” program (ANR-15-IDEX-02), grants from the Agence Nationale de la Recherche (ANR-15- CE11-0018-01), the Laboratory of Excellence GRAL financed within the Grenoble Alpes University graduate school (Ecoles Universitaires de Recherche) CBH-EUR-GS (ANR-17- EURE-0003), and the Fondation pour la Recherche Medicale (Team FRM 2017, DEQ20170336705). Julian Trouillon received a Ph.D. fellowship from the French Ministry of Education and Research. We further acknowledge support from CNRS, INSERM, CEA, and Grenoble Alpes University.

