## Supplementary Figures for "Determination of the Two-Component Systems regulatory network reveals core and accessory regulations across *Pseudomonas aeruginosa* lineages"

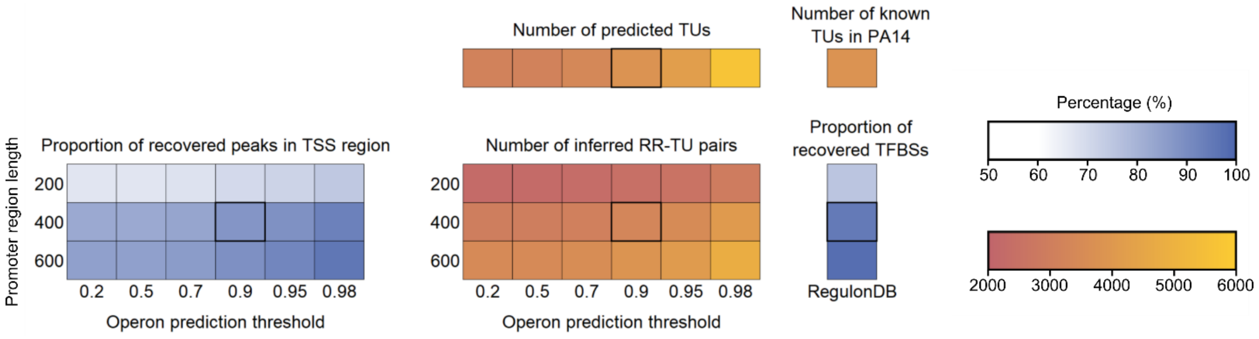


**Figure S1. Optimization of target inference parameters. Related to Figure 1.** 18 different combinations of promoter region length and operon prediction threshold were tested for TFBS recovery performance in comparison to experimentally obtained transcription start sites (TSS) in PA14 (Wurtzel et al., 2012) and known TFBSs relative position from RegulonDB (Santos-Zavaleta et al., 2019). The best-performing parameter pair was then chosen as allowing the highest recovery of known TFBS (>95%) while keeping the number of predicted TUs close to PA14 experimental number (0.994:1 ratio). The cells corresponding to the parameters chosen for the analysis are boxed in black. TU: Transcriptional Unit.

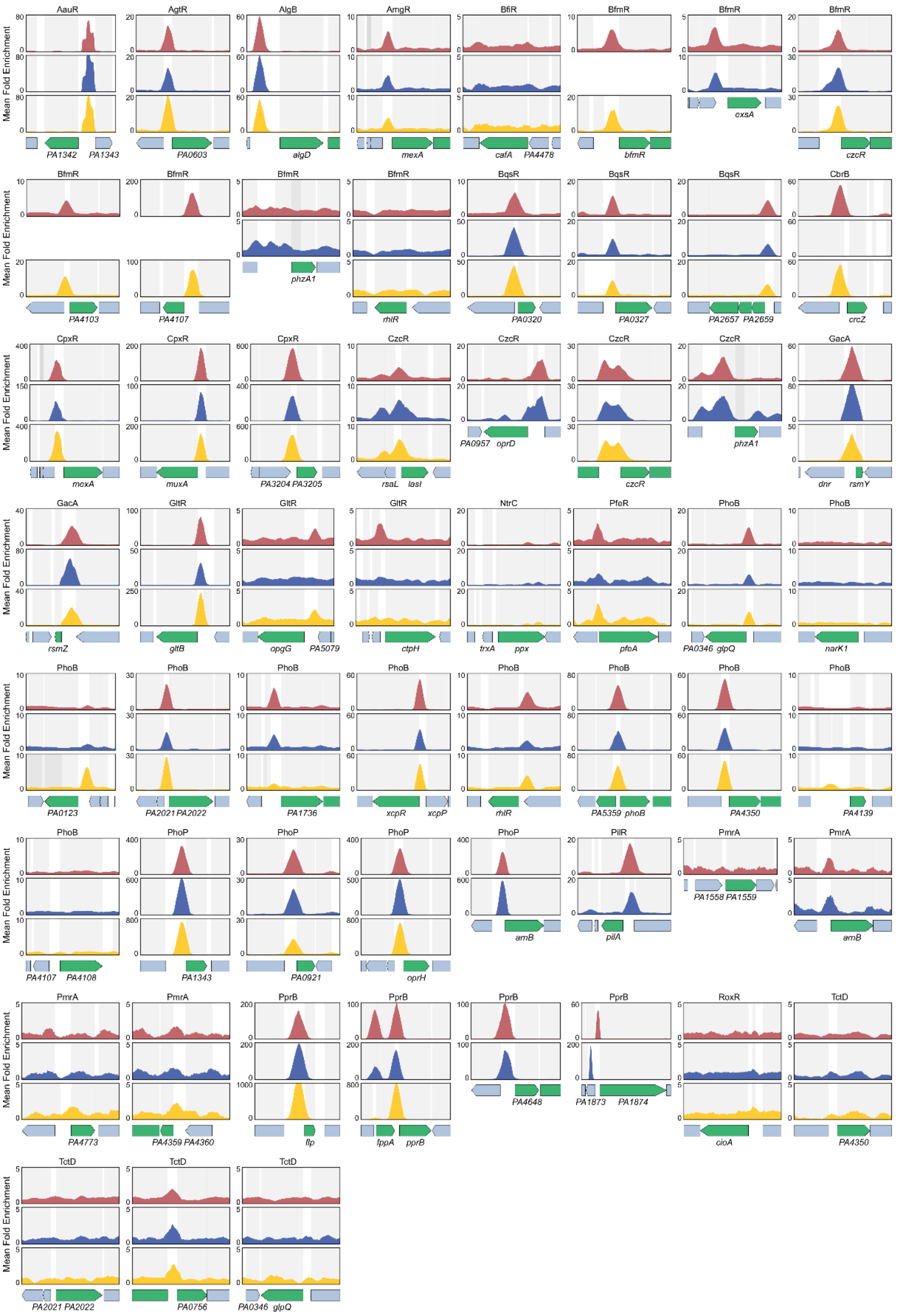


**Figure S2. Comparison with high-confidence known targets. Related to Figure 1.** For each high-confidence known target, enrichment coverage tracks of DAP-seq against negative controls are shown for the corresponding RR in all genomes (PAO1: red, PA14: blue, IHMA87: yellow) where the target gene is present. Known targets are shown as green arrows.

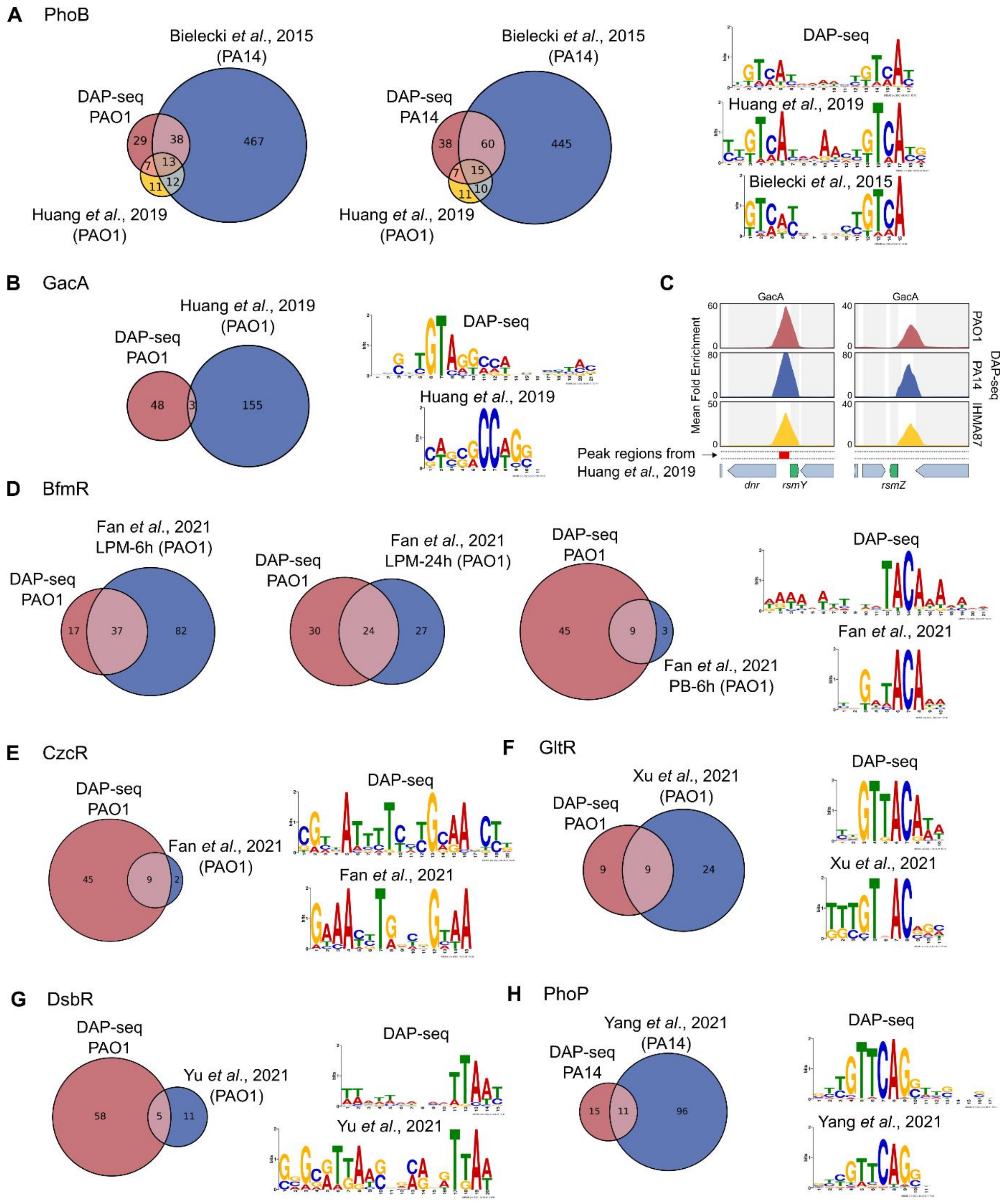


**Figure S3. Comparison of DAP-seq with published ChIP-seq results. Related to Figure 1.** Previously reported ChIP-seq experiments on PhoB (A), GacA (B), BfmR (D), CzcR (E), GltR (F), DsbR (G) and PhoP (H) were compared to the DAP-seq results. Target inference and DNA motif detection were performed in the same way between DAP-seq and ChIP-seq (Methods). (C) Enrichment coverage tracks of DAP-seq with GacA around *rsmY* and *rsmZ* genes. GacA binding regions reported in ChIP-seq by Huang *et al*. are shown as red boxes.

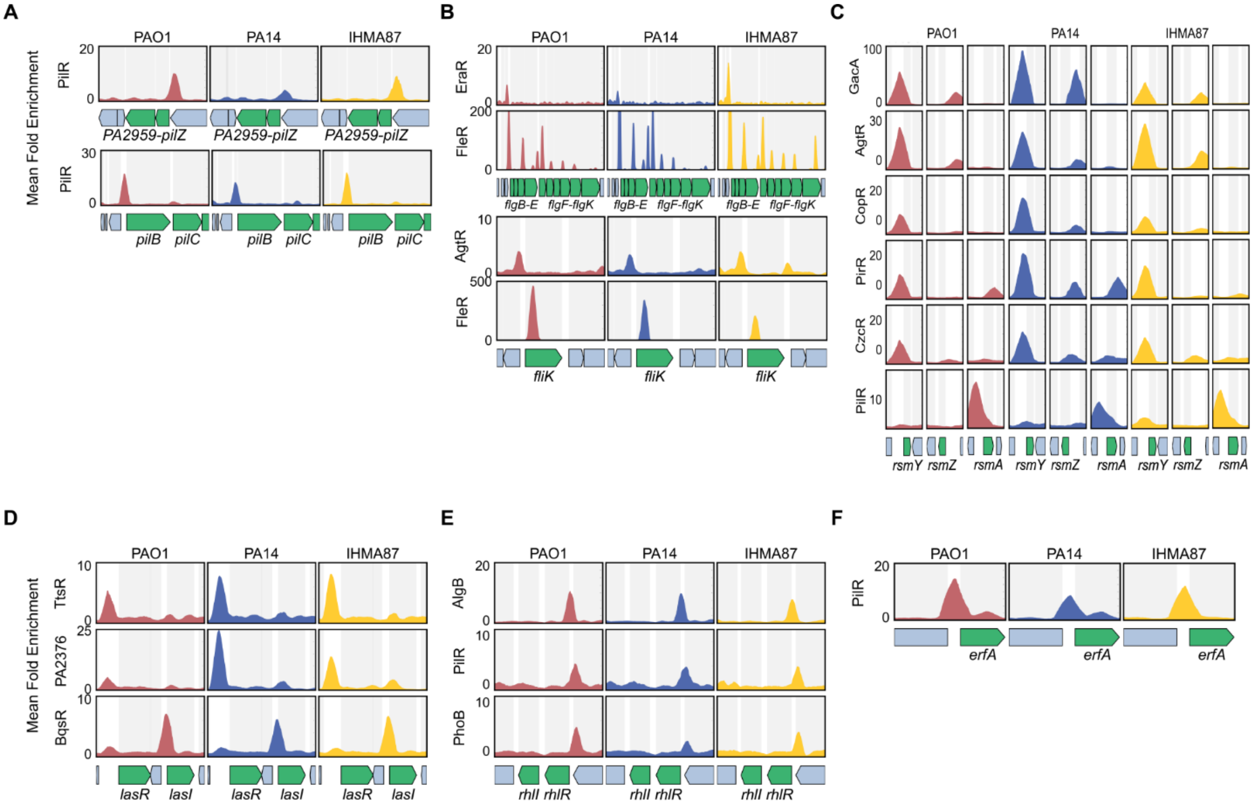


**Figure S4. The core TCS network is composed of major virulence-related targets. Related to figure 3.** Enrichment coverage tracks of DAP-seq against negative controls are shown for the corresponding RRs in all genomes (PAO1: red, PA14: blue, IHMA87: yellow) for *pilZ-PA2959* and *pilBC* (A), *flg* operons and *fliK* (B), *rsmY, rsmZ* and *rsmA* (C), *lasR* and *lasI* (D), *rhlRI* (E) and *erfA* (F). Inferred targets from the observed binding sites are shown in green.

**A**

| | Number of RRs binding | | |
|---|---|---|---|
| **Name** | **PAO1** | **PA14** | **IHMA** |
| *aprA* | 11 | 13 | 15 |
| *PA0123* | 7 | 16 | 15 |
| *xcpR-Z* | 13 | 10 | 9 |
| *flp* | 8 | 12 | 12 |
| *phzH* | 14 | 14 | 0 |
| *czcCBA* | 12 | 9 | 6 |
| *dppA3* | 9 | 9 | 9 |
| *nuoA-N* | 6 | 10 | 10 |
| *PA3601-PA3600* | 8 | 10 | 8 |
| *hupB* | 8 | 9 | 8 |
| *zipA* | 10 | 9 | 5 |
| *oprP* | 7 | 6 | 11 |
| *rsmY* | 8 | 7 | 8 |

**B**

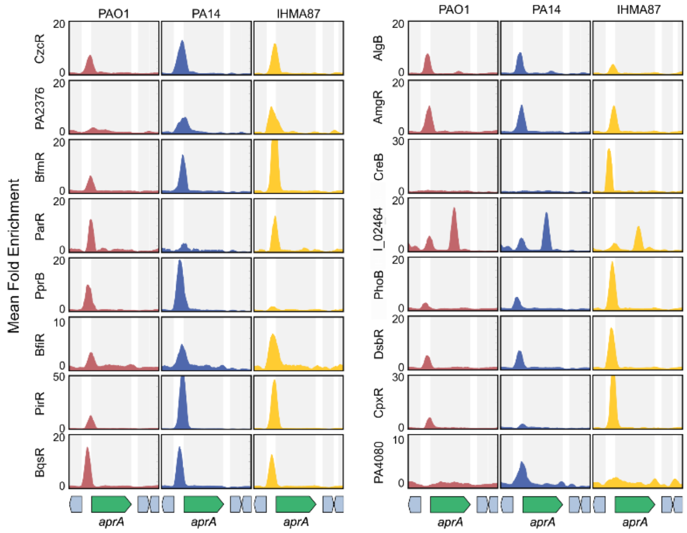


**Figure S5. The most targeted genes of the TCS regulatory network. Related to figure 4.** (A) Table of the top 10 most targeted genes on average. (B) Enrichment coverage tracks of DAP-seq against negative controls are shown for RRs binding to the *aprA* promoter region.
